# The bromodomain inhibitor JQ1 is a molecular glue targeting centromeres

**DOI:** 10.1101/2023.03.15.532673

**Authors:** Samuel Corless, Noor Pratap-Singh, Nezha S. Benabdallah, Jasmin Böhm, Alexander M. Simon, Vojtěch Dolejš, Simon Anders, Ana Banito, Sylvia Erhardt

## Abstract

Centromeres are the position on each chromosome that orchestrates the accurate partitioning of the genome during cell division. Centromere-dependent cell-cycle checkpoints are maintained by cancer cells to prevent catastrophic chromosome segregation defects in dividing cells^1, 2^, making centromeric chromatin a valuable target for anti-cancer therapeutics. However, no compounds have been identified that specifically target centromeric chromatin using standard drug discovery approaches. Here we develop a big-data approach to identify the protein composition of repetitive DNA loci, including centromeres, and screen candidate small molecules that act on centromeric chromatin composition. We discover that the BET bromodomain protein BRD4 localises to centromeres and regulates centromeric cohesion. We further show that the bromodomain inhibitor JQ1 affects centromeric BRD4 by stabilising a direct interaction between BRD4 and Centromere Protein B (CENP-B), acting as a molecular-glue that promotes centromere cohesion in a CENP-B-dependent manner. Strikingly, CENP-B transitions from a non-essential protein in JQ1-sensitive cells to the most significant determinant of cell-proliferation in JQ1-resistant cells. Our observations demonstrate a completely overlooked role for BRD4 and JQ1 in directly targeting the centromere, with important consequences for JQ1-derivatives currently entering clinical use^3^.

---

Human centromeres have long been considered an enigmatic and challenging region of the genome to study using sequencing-based approaches, due to the highly repetitive nature of their DNA (Fig. 1a)^4^. The function of centromeric loci is to ensure the accurate partitioning of the genome during cell-division by assembling the multi-subunit kinetochore complex, which attaches chromosomes to the spindle microtubules^5, 6^. Defects in centromere structure and function are key drivers of aneuploidy, tumour heterogeneity and cancer progression^7, 8^. Such diverse roles indicate that centromeric proteins are an attractive target for anti-cancer therapeutics, but there are no known inhibitors that specifically target components of centromeric chromatin. A major barrier to progress in human centromere biology has been the lack of studies mapping chromatin proteins to highly-repetitive DNA regions^9^, and for the most-part our understanding of centromeric chromatin remains in the pre-genomic era.

**Fig. 1.**
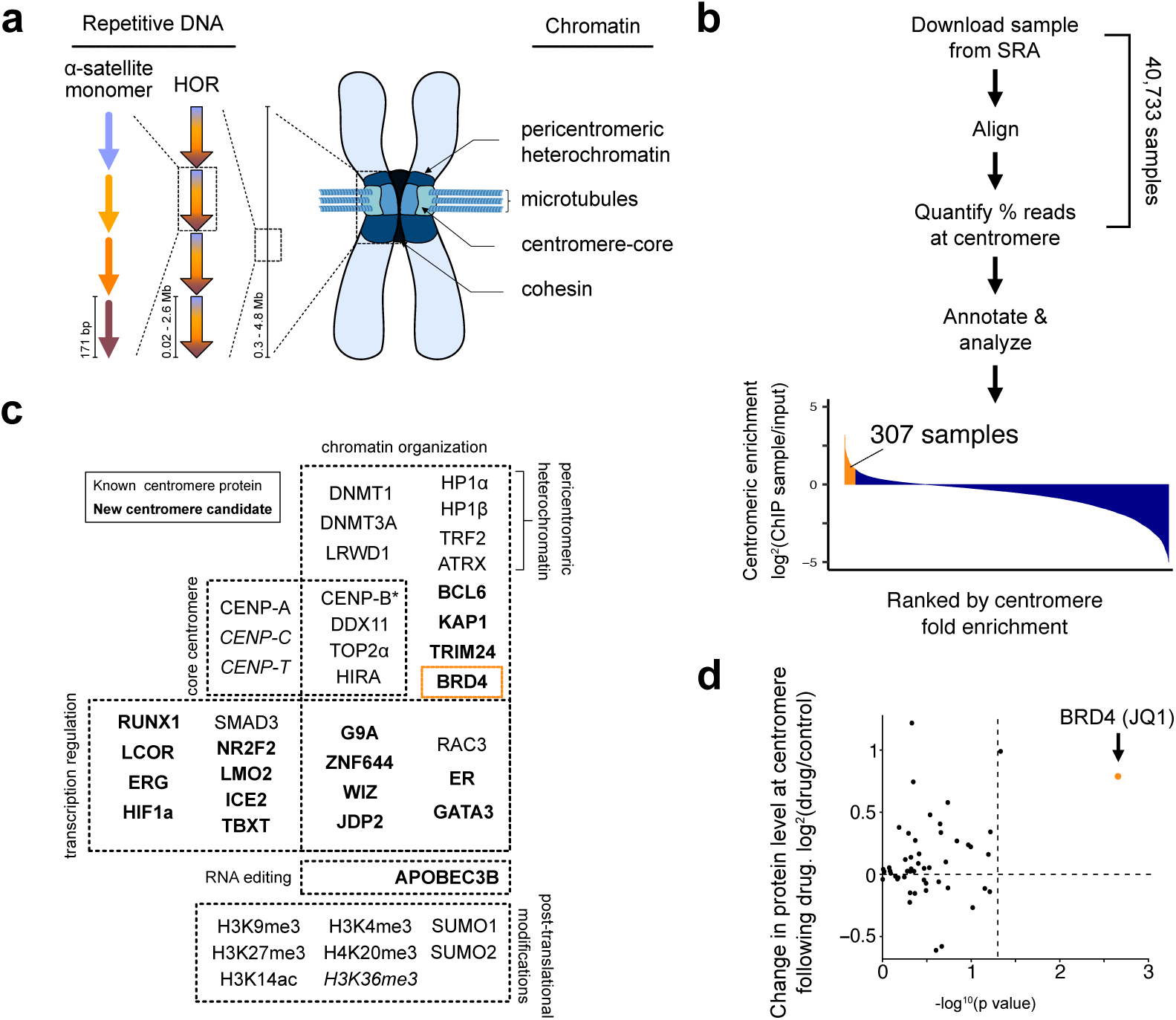
BRD4 is a centromere protein which is further enriched by the bromodomain inhibitor JQ1. **a**, Overview of human centromeric DNA organisation and the subdomains of centromeric chromatin displayed on metaphase chromosomes. Length of DNA repeat motifs given in base-pairs (bp) and Megabases (Mb). **b**, Summary of the REDPOINT workflow for centromeric loci. **c**, REDPOINT identifies 35 high confidence centromere proteins including known and novel candidates. Protein categories were determined from published data and Uniprot information. *CENP-B was identified using the same pipeline to analyse a published CUT&RUN dataset PRJNA546288. **d**, Focused analysis of 50 different protein-inhibitor pairs investigated in multiple studies and identified through the REDPOINT approach. The x-axis is the statistical significance from a paired-sample t-test comparing centromeric protein abundance in inhibitor treatment to control across multiple studies, with the vertical dotted line signifying a p-value of 0.05. The y-axis is the fold-change in protein abundance at the centromere following inhibitor treatment, with a positive fold change indicating more protein at the centromere and a negative fold change indicating less protein at the centromere.

In contrast to centromeres, our understanding of chromatin within unique-sequence regions has been revolutionised thanks to the broad application of genome-wide chromatin mapping approaches, which have significantly improved our understanding of the molecular mechanisms of transcription factors^10^, regulatory elements^11^, 3D genome organisation^12^, chromatin-associated disease^13, 14^ and specific classes of small-molecule inhibitors targeting chromatin components^13^. In these published studies centromeric loci were excluded from the analysis, either through the use of a genome-build in which centromeric repeats are absent or because standard pipelines exclude repetitive DNA^4^, but the raw experimental data containing this information is publicly accessible through the Sequence Read Archive (SRA)^15^. We took advantage of this vast resource and developed an approach called Reanalysis of Every Dataset at Positions Of INTerest (REDPOINT) that utilised published chromatin-mapping datasets to identify the protein composition of any locus. The objective of our study was to identify inhibitors that target the protein composition of centromeres. To achieve this, we first analysed the proteins present at centromeric chromatin and then conducted a focused investigation on a subset of REDPOINT data that included small-molecule treatments.

Our analysis included 40,733 chromatin immunoprecipitation sequencing (ChIP-seq) experiments that identified the genomic distribution of more than 1200 human chromatin proteins and post-translational modifications (Fig. 1b). The workflow included the following steps: 1) download 100,000 reads per ChIP-seq sample from the SRA, 2) align to a human genome build which includes information for repetitive loci^4^, 3) retain repetitive regions and identify the percentage of reads aligned to the centromere compared to the whole genome, 4) repeat for each dataset, 5) annotate and analyse the data. We observe a striking enrichment of proteins previously identified at the centromere-core which is the site of kinetochore formation and microtubule attachment during chromosome segregation^16^ (CENP-A, CENP-B, CENP-C, CENP-T), pericentromeric heterochromatin which supports centromere-core function^17^ (HP1α, HP1β, DNMT1, DNMT3A, ATRX) and other proteins with centromeric roles (LRWD1, TRF2, DDX11, TOP2α, HIRA, RAC3, SMAD3, BCL6) (Fig. 1a, c). We also identified 19 high confidence new centromere protein candidates, which met one or both of the following criteria: 1) more than two-fold enriched compared to corresponding control, 2) in the top 500 samples by percent of reads aligned to the centromere. Within the new centromere candidates was BRD4, a key protein in the organization of super-enhancers, transcription elongation and interphase genome organization^18–20^.

By analysing a particular subset of the REDPOINT data that contained ChIP-seq samples acquired post-exposure to diverse small-molecule inhibitors, and focusing specifically on 50 protein-inhibitor pairs that were examined in multiple studies, we identified that BRD4-JQ1 is the sole protein-inhibitor pairing that exhibits a highly significant alteration in protein abundance at the centromeric regions across all studies that were analysed (Fig. 1d). JQ1 is a small-molecule BET bromodomain inhibitor that prevents BRD4 from binding its acetyl-chromatin substrate^21^, so the canonical mechanism-of-action predicts that JQ1 treatment decreases BRD4 levels at its chromatin binding sites. Unexpectedly, we observe that JQ1 treatment consistently increases BRD4 occupancy at centromeres.

The discovery of the presence and JQ1-induced increase of BRD4 at centromeres is remarkable. Typically, BRD4 is considered as a regulator of genes and enhancers^19^, but centromeric regions lack these elements and are mostly transcriptionally inactive^9, 22^. Recent studies have revealed that BRD4 also plays a role in the organisation of the interphase genome by promoting cohesin loading at TAD boundaries, indicating a role in contemporary theories of genome organisation through chromatin loop extrusion^12, 18, 23^. However, a link between BRD4 and centromeric cohesion has not previously been proposed, which led us to investigate the role of BRD4 and the inhibitor JQ1 at centromeres.

## BRD4 regulates centromere cohesion

To visualise the relationship between BRD4 and centromeres in greater detail we mapped BRD4 ChIP-seq reads from diverse cell types onto the telomere-to-telomere (T2T) build of chromosome 8^24^. BRD4 is enriched at many sites, including at centromeric loci defined by the co-localisation with CENP-A and HP1 (Fig. 2a). Immunofluorescence in HeLa cells with antibodies targeting BRD4 and CENP-A revealed a clear co-localisation of these proteins as puncta within the DAPI dense heterochromatin at the periphery of nucleoli (Fig. 2b), indicating that BRD4 is a centromere-core protein not enriched in adjacent pericentromeric heterochromatin.

**Fig. 2.**
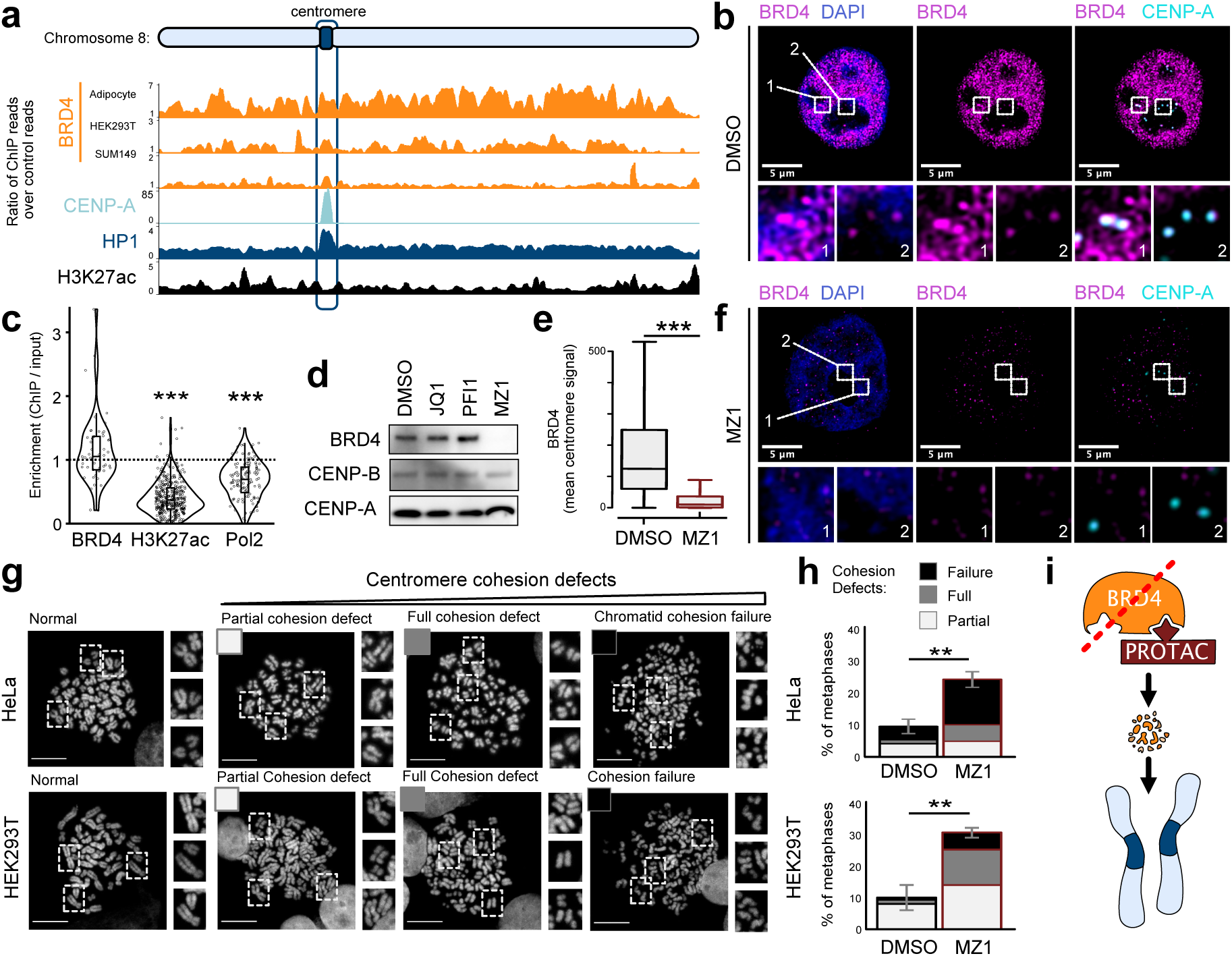
BRD4 degradation disrupts mitotic centromere cohesion. **a**, Telomere-to-telomere distribution of BRD4, CENP-A, HP1 and H3K27ac on chromosome 8. Data displays the ratio of ChIP reads compared to a matched input from the same study. Datasets: CENP-A (HeLa) PRJNA183936, BRD4 and HP1 (HEK293) PRJNA171165, H3K27ac and BRD4 (SUM149) PRJNA268319, BRD4 (adipocyte) PRJNA270506. **b**, Immunofluorescence imaging of BRD4 (magenta) and CENP-A (cyan) in HeLa cells treated for 24 hours with DMSO. **c**, Centromere enrichment of all BRD4, H3K27ac and RNA polymerase II (Pol2) ChIP-seq samples in the REDPOINT dataset compared to the corresponding control from the same study. Statistical analysis t-test between BRD4 and H3K27ac/Pol2 (*** p<0.001). **d**, Western blot analysis of BRD4, CENP-A and CENP-B following 24 hours treatment with two bromodomain inhibitors ((±)-JQ1 and PFI1) and the BRD4-specific PROTAC MZ1. **e**, Quantification of BRD4 signal intensity at the core-centromere in HeLa cells following 24 hours DMSO or MZ1 treatment. Values are the intensity of BRD4 signal overlapping with foci of CENP-A. DMSO (n=3707 centromeres), MZ1 (n=3505 centromeres). Whiskers = boxplot standard in R (1.5x IQR). Statistical analysis t-test (*** p<0.001). **f**, Immunofluorescence imaging of BRD4 (magenta) and CENP-A (cyan) in HeLa cells treat following 24 hours MZ1 treatment. Brightness/contrast settings the same as Fig. 1b. g, Representative mitotic chromosomes from HeLa and HEK293T cells with normal morphology and increasing severity of centromere cohesion defect. Identifier for subsequent quantifications in inset boxes. **h**, Quantification of mitotic chromosome defects following 24 hours MZ1 treatment in HEK293T and HeLa cells. Error bars = standard error of the mean. Statistical analysis based on triplicate experiments, t-test (** p < 0.01). **i**, Model outlining the effect of BRD4 degradation on mitotic chromosome structure.

Centromeres have unique chromatin properties that differ markedly from gene-rich regions, including low levels of histone acetylation and transcription^22, 25, 26^, indicating a non-canonical role for BRD4 at centromeric chromatin. We first confirmed that centromeric BRD4 enrichment does not correspond to an enrichment in H3K27ac or RNA polymerase II (Fig. 2a, c). Next we completely depleted BRD4 from cells using MZ1, a proteolysis-targeted chimera (PROTAC) small-molecule that selectively degrades BRD4 when compared to BET bromodomains paralogs BRD2 and BRD3^27^. Unlike the bromodomain inhibitors PFI1 and the racemic mixture of (±)-JQ1 (referred to as JQ1), MZ1 treatment leads to the complete loss of BRD4 protein from the cell (Fig. 2d), including at the centromere core (Fig. 2e, f). We then quantified the effect of BRD4 degradation on centromere cohesion defects in metaphase chromosomes prepared from HEK293T and Hela cells, using a scale ranging from partial centromere defect to complete cohesion failure (Fig. 2g). This revealed that MZ1 treatment led to a highly significant increase in centromere cohesion defects (Fig. 2h). These results identify a previously uncharacterised role for BRD4 in the organisation of centromere cohesion and demonstrate that PROTAC inhibitors targeting BRD4 strongly disrupt metaphase chromosome architecture in different cell types (Fig. 2i).

## JQ1 promotes BRD4 at centromeres

To understand why JQ1 promotes BRD4 localisation to centromeres (Fig. 1d) we investigated in more detail data from 15 different studies with 26 different cell-lines (Fig. 3a, b). In all studies and in 25 of 26 cell-lines BRD4 increased at the centromere following JQ1, with the JQ1-resistant cell-line SUM149R being the only exception. This recruitment was specific to BRD4 and to centromeres, since other bromodomain proteins did not show the same level of centromere accumulation (Extended Data Fig. 1a) and BRD4 was not enriched at abundant LINE2 repetitive DNA elements following JQ1 treatment (Extended Data Fig. 1a, b). To confirm this specificity, we plotted the fold-change in BRD4 localisation following JQ1 treatment on a T2T genome and observed that the centromere is the principle locus gaining BRD4 association in different cell lines (Fig. 3c). Importantly, JQ1 treatment did not significantly increase H3K27ac, RNA polymerase II or centromeric transcription ruling out a JQ1-dependent switch to the canonical functions of acetylation-dependent transcriptional control at centromeres (Fig. 3c, d, Extended Data Fig. 1a, c).

**Fig. 3.**
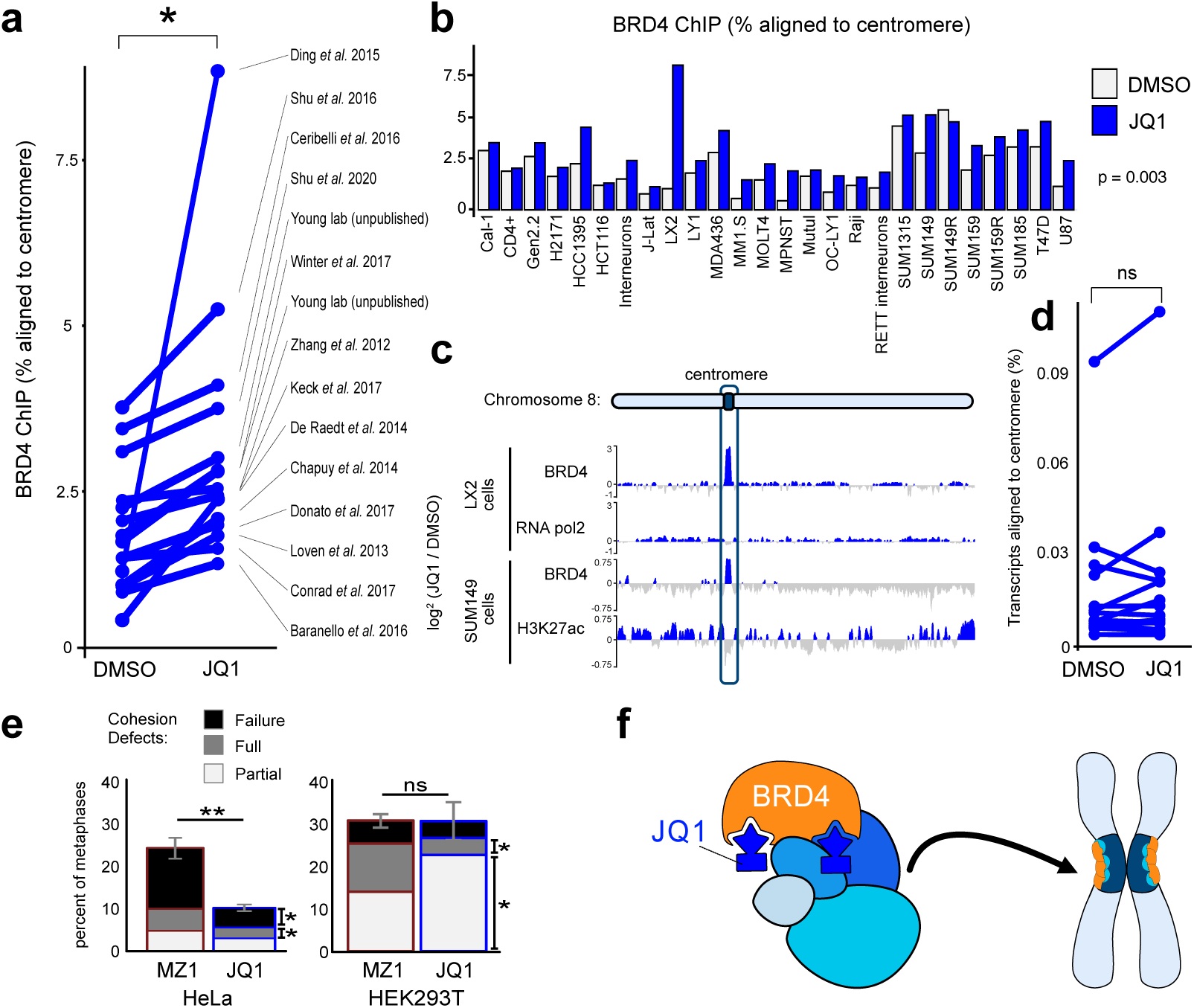
JQ1 promotes a non-canonical cohesion function of BRD4 at centromeres. **a**, Analysis of 15 different ChIP-seq datasets from studies comparing BRD4 distribution in control and JQ1 treated cells. Paired t-test p value < 0.05. **b**, BRD4 is more enriched following JQ1 treatment in 25 out of 26 analysed cell lines. Statistical analysis paired-sample t-test (p=0.003). **c**, BRD4, RNA polymerase II and H3K27ac enrichment following JQ1 compared to DMSO treatment aligned to the T2T genome of chromosome 8. LX2 data from PRJNA253142, SUM149 data from PRJNA542646. **d**, Meta-analysis of RNA-seq datasets comparing DMSO and JQ1 treatment from 12 studies. **e**, Quantification of mitotic chromosome defects following 24 hours MZ1 or (±)-JQ1 treatment in control HEK293T and HeLa cells. Indicator as outlined in Fig. 2g. Error bars = standard error of the mean. Statistical analysis based on triplicate experiments, t-test (ns ‘not significant’, * p<0.05, ** p<0.01). **f**, Model outlining the rescue of centromere cohesion defects in JQ1-treated cells that recruit BRD4 to the centromere.

To consider non-canonical effects of JQ1 treatment we again quantified centromere cohesion defects in metaphase chromosomes prepared from HEK293T and Hela cells. In contrast to PROTAC-based degradation, JQ1 treatment identified fewer centromere cohesion defects (Fig. 3e). HEK293T cells show a significant transition towards milder defects (‘mixed’ phenotype). More strikingly, the proportion of HeLa cells with chromosome defects following JQ1 treatment was comparable to DMSO (Fig. 2h, 3e), suggesting the possibility that BRD4 recruitment stabilises centromeric chromatid cohesion (Fig. 3f).

## JQ1 stabilises an interaction between BRD4 and CENP-B

To further characterise the effect of JQ1 treatment on BRD4 at centromeres we performed immunofluorescence and observed no large-scale reorientation of BRD4 towards centromeric loci (Extended Data Fig. 2a, b), reflecting the subtle yet significant change in BRD4 observed by REDPOINT (Fig. 3b) and highlighting the higher sensitivity of our approach to determine novel chromatin interactions than can be achieved by microscopy-based screens.

To investigate at the molecular scale how BRD4 is further recruited to and maintained at the centromere upon JQ1 treatment we analysed published co-IP mass spectrometry (coIP-MS) data identifying the proteins which specifically interact with BRD4 following JQ1 treatment^28, 29^. JQ1 treatment consistently induced a strong, and previously overlooked, association between BRD4 and key centromere proteins (Fig. 4a, Extended Data Fig. 2c, d). Across all four cell-types assayed, and in as little as 10 minutes following JQ1 addition, a new top interaction between BRD4 and the constitutive centromere protein CENP-B formed (Fig. 4a, Extended Data Fig. 2c, d). Co-IP MS for other BET proteins show a low-affinity (BRD3) or no (BRD2) interaction with CENP-B following JQ1 (Extended Data Fig 2e). Members of the MIS18 complex (MIS18A, MIS18B and MIS18BP1), which binds the centromere for a few minutes each anaphase to licence the deposition of CENP-A^30^, were also highly enriched in coIP-MS experiments but considered unlikely to be the major factor recruiting BRD4 to the centromere due to the transient nature of this interaction.

**Fig. 4.**
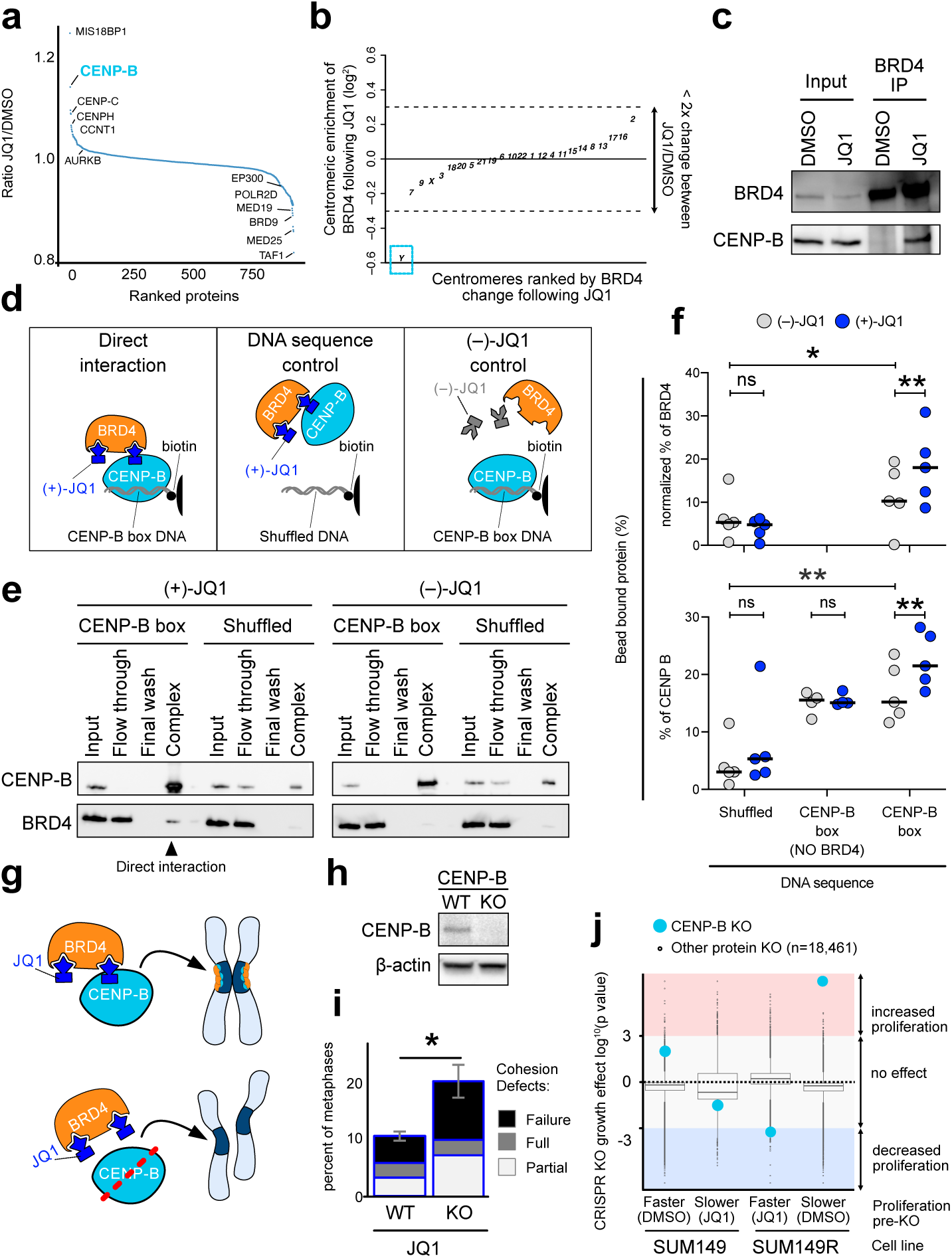
JQ1 is a molecular glue that retains BRD4 at centromeres through CENP-B. **a**, Co-IP MS identifying the change in proteins interacting with BRD4 following JQ1 treatment, with key proteins for centromere identity and BRD4-mediated transcription regulation highlighted. Data from Shu et al. (2020)^28^. **b**, ChIP-seq analysis of centromeres ranked by BRD4 enrichment following JQ1 treatment in male cells. Enrichment was calculated by identifying the percentage of centromeric BRD4 at each chromosome in a sample, and then calculating the ratio of JQ1 treated sample compared to DMSO treated sample. The dotted lines indicate a 2x change in the abundance of a BRD4 at a centromere, the blue square highlights the Y chromosome. Data from: PRJNA253142. **c**, Co-IP for BRD4 following DMSO or (±)-JQ1 treatment for 1 hour in HeLa cells, followed by western blot for BRD4 and CENP-B. **d**, Model outlining an *in vitro* reconstitution pull-down experiment to determine whether a direct interaction occurs between BRD4^bromo^, (+)-JQ1 and CENP-B, with control experiments containing (–)-JQ1 or shuffled DNA sequence. **e**, Example *in vitro* reconstitution pull-down experiment result comprised of western blots of BRD4 and CENP-B. BRD4 is at 10x molar excess compared to CENP-B. ‘Input’ is 10% of the total sample and ‘flow through’/’wash’ is 10% of the total. ‘complex’ is 100% of the bead bound protein. **f**, Quantification of independent *in vitro* reconstitution pull-down experiments. To account for the ten-fold molar excess of BRD4 the ‘Bead bound protein (% input)’ for BRD4 was multiplied by ten to give the ‘Normalized bead bound protein (%)’. Statistical tests are paired-sample t-test (* p<0.05, ** p<0.01). **g**, Model outlining the predicted outcome of CENP-B loss on JQ1-linked centromere cohesion. **h**, Western blot of control (WT) and CENP-B KO HeLa cells probed with anti-CENP-B antibodies and β-actin as loading control. **i**, Quantification of cohesion defects in 24 hours (±)-JQ1 treated HeLa control and CENP-B KO cells. Error bars = standard error of the mean. Statistical analysis based on triplicate experiments t-test (* p<0.05). **j**, CRISPR KO screen comparing the effect of CENP-B KO relative to 18,460 other proteins on cellular proliferation in JQ1-sensitive and -resistance SUM149 breast cancer cells. Data from Shu et al. (2020).

CENP-B is the only well characterised centromere-specific DNA binding protein, and binds throughout the cell-cycle to a 17 base-pair CENP-B box DNA sequence found on every centromere except the Y-chromosome^31^. We observe a strong positive correlation between the level of CENP-B binding at a centromere and the recruitment of BRD4 following JQ1 treatment (Extended Data Fig. 3a). In fact, JQ1 treatment recruits BRD4 to all centromeres except the Y chromosome (4 out of 5 male cell-lines investigated) (Fig. 4b, Extended Data Fig. 3b, c). No correlation was found between CENP-B and RNA polymerase II following JQ1 treatment, or BRD4 following degradation with the PROTAC inhibitor dBET6 (Extended Data Fig. 3d). Together, these observations strongly support CENP-B as the key protein mediating the JQ1-dependent recruitment of BRD4 to centromeres.

We independently confirmed a strong and specific JQ1-dependent interaction between BRD4 and CENP-B in HeLa and HEK293T cells by co-IP western blot (Fig. 4c, Extended Data Fig. 4a, b), which was not observed in co-IP following treatment with the negative enantiomer (–)-JQ1^32^ or the BET bromodomain inhibitor PFI1^33^ which has a different chemical structure (Extended Data Fig. 4c).

To identify whether the JQ1-dependent interplay between BRD4 and CENP-B is direct, we reconstituted the interaction *in vitro* using full-length CENP-B and the bromodomains of BRD4 (BRD4^bromo^) (Fig. 4d). We observed that CENP-B protein was strongly associated with the CENP-B box DNA sequence when compared to a shuffled version of the same sequence (Fig. 4 e, f). The addition of (+)-JQ1 or (–)-JQ1 enantiomer to CENP-B alone did not change the level of interaction with the CENP-B box DNA sequence, indicating that these small-molecules have no effect on CENP-B-DNA interactions alone. BRD4^bromo^ was enriched in the presence of CENP-B bound CENP-B box DNA compared to shuffled DNA, even with the (–)-JQ1 enantiomer, indicating that a low-affinity complex between BRD4 and CENP-B occurs independently of the BRD4-JQ1 association *in vitro*. However, with the addition of the (+)-JQ1 enantiomer, BRD4^bromo^ and CENP-B became significantly more enriched, indicating that (+)-JQ1 stabilises the BRD4-JQ1-CENP-B-DNA complex *in vitro*. The stabilisation of a low-affinity interaction is one of the defining properties of a rare class of inhibitors called molecular-glues, which enable gain-of-function interactions that can target previously undruggable pathways^34, 35^, and our evidence suggests that (+)-JQ1 is the first centromere-targeting small-molecule via this mechanism-of-action.

## BRD4-JQ1-CENP-B stabilises centromere cohesion

If the BRD4-JQ1-CENP-B complex stabilises the centromere cohesion role of BRD4, removing CENP-B and treating cells with JQ1 should give centromere cohesion defects comparable to MZ1 treatment (Fig. 4g). To test this hypothesis, we generated HEK293T and HeLa cells lacking CENP-B through a CRISPR knock-out (CENP-B KO) approach (Fig. 4h, Extended Data Fig. 5a). In HEK293T CENP-B KO cells we observed a marginal enhancement of the already apparent cohesion defect following JQ1 treatment (Extended Data Fig. 5b). However, in HeLa CENP-B KO cells we observed a massive increase in centromere cohesion defects compared to control cells, comparable to the degradation of BRD4 and consistent with our proposed model (Fig. 4i). These observations indicate that JQ1 has a cell-type variable role in targeting centromere structure and function, through a novel complex between CENP-B and BRD4.

Interestingly, among the 26 different cell lines analysed in this study, only the JQ1-resistant cell-line SUM149R^28, 36^ did not show an increase in centromere localised BRD4 following JQ1 treatment (Fig. 3b). SUM149R cells were selected in increasing concentrations of JQ1 and now grow more rapidly in the presence of JQ1 than in DMSO^36^. To determine whether the mechanism of resistance is SUM149R cells is linked to CENP-B, we analysed a published CRISPR-KO screen in parental and resistant cells treated with DMSO or JQ1^28^. CENP-B is a non-essential protein in SUM149 (wild-type) cells treated with DMSO or JQ1 because knock-out does not significantly alter cellular proliferation, consistent with published observations including viable CENP-B KO mice^37^. In contrast, SUM149R (JQ1 resistant) cells are highly sensitive to the loss of CENP-B protein (Fig. 4j). In rapidly growing SUM149R-JQ1 cells, CENP-B KO reduces cell proliferation, whereas in slowly growing SUM149R-DMSO cells, CENP-B KO gives the most significant increase in cell proliferation out of 18,461 proteins tested. We identified no other protein with a similar response amongst BET bromodomain proteins, transcription elongation components, cohesin complex components or components of centromeric chromatin (Extended Data Fig. 5c). These observations suggest that CENP-B specifically regulates cell proliferation in a cellular model of JQ1 resistance and indicates that the BRD4-JQ1-CENP-B complex is critical in establishing JQ1 resistance.

## Discussion

In summary, our study identified a previously overlooked role for BRD4 and the bromodomain inhibitor JQ1 in centromere cohesion. We demonstrated JQ1 as the first small-molecule directly targeting centromeric chromatin, through a molecular-glue mechanism-of-action that stabilises a link between BRD4 and CENP-B. The identification of molecular glues is rare, but each new example opens up the possibility of targeting previously undruggable pathways^34, 35^. Future optimisation of JQ1 structure could form the basis of a new class of centromere drugs targeting chromosome segregation, a process which is important only in dividing cells. We showed that the BRD4-JQ1-CENP-B interaction stabilises centromere cohesion in mitosis, supporting a direct role for BRD4 in centromere cohesion and complementing recent observations that identified a role for BRD4 in regulating TADs by acting upstream of cohesin loading^18, 38^. Finally, we show that CENP-B transitions from non-essential in JQ1-sensitive cells to the most significant determinant of cell proliferation in JQ1-resistant cells. A significant challenge in employing BET bromodomain inhibitors for cancer treatment is the swift emergence of resistance^3, 39^, and our observations show that this resistance can occur via mechanisms independent of gene-regulation. Future bromodomain inhibitors developed for clinical use must consider this centromere role, in order to better target cancer-cell pathways and avoid mechanisms of resistance linked to CENP-B.

## Methods

### Candidate identification with REDPOINT

#### Dataset curation for analysis by REDPOINT

A list of human ChIP-seq accession numbers, which allow dataset recovery from the Sequence Read Archive (SRA)^1^, were obtained using the ‘advanced search’ feature of the European Nucleotide Archive (ENA) (https://www.ebi.ac.uk/ena/browser/advanced-search) with the search parameters: Data type = ‘Raw reads’, Taxon name = ‘Homo sapiens’, Library Strategy = ‘ChIP-seq’, and the field parameters: ‘experiment accession’ and ‘sample accession’. The ‘sample accession’ enabled each dataset to be downloaded from the SRA, and the ‘experiment accession’ allowed the samples from the same study to be grouped in downstream analyses. The state of sample annotation within the SRA, ENA or the Gene Expression Omnibus (GEO) did not allow for automated annotation of key features for our study, including whether the sample was from a ChIP-seq experiment or corresponding input/control, the protein enriched by the ChIP-seq experiment, the tissue/cell-line investigated, whether the sample was treated with a drug or other experimental perturbation, and the associated PubMed ID. To enable detailed downstream analysis of protein enrichment relative to corresponding input/control, the influence of cell/tissue type, and the effect of perturbations including drug treatment, the first 21,462 samples from 1,234 studies were annotated by hand using the relevant parameters included with the samples GEO submission.

#### Genomic positions investigated by REDPOINT

We used the hg38 genome build, which contains the entire human genome including representative maps of repetitive regions, from which we extracted centromeric and LINE2 alignment information^2^.

#### Detailed description of centromeric loci

Centromeric repeats, are annotated in the hg38 genome and in recently published T2T genomes^2–4^. In hg38 centromeric DNA is divided into different classes of satellite repeat, including the dominant α-satellite repeat found at the centromere on every chromosome and low abundance satellite classes which are each found at a small number of centromeres (GSAT, GSATII, GSATX, HSAT4, SST1). The CENP-A-containing core centromere has only been shown to form on the dominant α-satellite DNA^3^. α-satellite is highly divergent and has different Higher Order Repeats (HORs) on different chromosomes^3^. In hg38 the centromeric α-satellite repeats are sub-divided into different chromosomes based on the abundance of different HORs, which allows the chromosome-specific abundance of centromeric proteins to be determined from ChIP-seq data and demonstrates the detailed nature of the centromeric annotation in this genome build. Recently, telomere-to-telomere (T2T) builds of the human genome, with the centromeric sequence in the correct linear order across the entire domain for an individual pseudo-haploid genome, have been published^3, 4^. Between cell-lines/chromosomes/individuals there is a strong divergence in the position and abundance of different α-satellite HORs, as well as different epi-allele positioning of key centromere proteins including CENP-A within the α-satellite DNA, and mapping short-read ChIP-seq data from a diversity of sources to the complete T2T genome provides little extra positional information compared to the hg38 genome. We therefore identified candidate proteins using the hg38 genome, some of which we validate by aligning to a T2T genome.

#### REDPOINT analysis

The REDPOINT pipeline utilises standard bioinformatic tools for data acquisition, genome alignment and genomic position investigation. Briefly, 1.) For each sample, as determined by the SRA ‘sample accession’, we downloaded a sub-set of 100,000 reads using ‘fastq-dump’ from NCBI SRA Tools (parallel-fastq-dump -sra-id SAMPLE_ID --threads 4 --outdir out/ --split −3 --gzip). The −3 split argument automatically differentiates between single-end and paired-end sequencing. For consistency across the analysis, and because our analysis focuses on highly repetitive DNA, we only utilised the first read of paired-end sequencing in downstream analysis. We selected reads starting at read 10,000 to avoid observed problems at the start of a file and obtain a random assortment of reads (https://edwards.flinders.edu.au/fastq-dump/). In all cases where we compared a sample sub-set aligned to the complete sequencing run we observed an exact correspondence in the percentage of reads at the centromere, indicating that 100,000 reads are sufficient to identify protein enrichment for a locus comprising 2.15% of the genome without generating sampling bias. 2) We used Bowtie2^5^ (ver.2.3.0) to align reads to the hg38 build of the human genome using default settings. 3) We identified the percentage of reads at each genomic position (e.g. centromeric loci, ACTB gene promoter, etc.) using HTseq^6^. This information was recorded and then the loop re-set to begin the next ChIP-seq dataset.

#### Quality control of ranked protein enrichment

The first method utilised for identifying the proteins associated with different loci of interest was to rank samples based on the percentage of ChIP-seq reads associated with each locus. This allowed the use of the full 40,734 samples analysed in the REDPOINT approach and identified well characterised proteins associated with centromeric chromatin. We also noted many examples of artefact proteins enriched via this approach. For example, there were samples which were very highly enriched at centromeric loci, comparable to key CENP proteins, but upon further inspection showed no enrichment compared to control samples within the same study – indicative of a problem with the entire study rather than a new centromeric protein. This suggested to us that the ‘ranked data’ approach was sufficient to identify candidate proteins, but could be improved upon by comparing samples to corresponding input/control.

#### Quality control of fold-enriched proteins

The second method for identifying the proteins associated with different loci of interest had a number of quality control steps which take advantage of the 21,462 annotated ChIP-seq samples. Input and control samples provided an invaluable quality control metric, firstly to exclude problematic datasets and secondly to benchmark the proportion of centromeric DNA identified in the samples within a study, which varied by around four-fold from around 1% to around 4% for the majority of samples. Analysis of genomic DNA samples in the same cell line across multiple studies suggested that these differences reflects inter-study variability, rather than biological variability, which may be linked to differential PCR efficiency of highly repetitive loci or variability in the loss of highly compact centromeric chromatin dependent on the methodology used for genomic DNA or control preparation^7^. Many studies did not include a genomic input sample, IgG pull-down or other corresponding control and we excluded these, retaining 11,881 samples from 887 studies. Focussing on the genomic DNA or control samples in these studies, we noted that a sub-set of these samples contained virtually no DNA which aligned to centromeric loci. This is not biologically possible and suggests a technical problem with these samples, which may reflect experimental issues or mistakes in data handling. We removed all studies with fewer than 1% of the reads aligned to centromeric DNA which left us with 9,764 samples from 780 studies, i.e. 63% of the annotated studies. For each of these samples we calculated the fold-enrichment for each ChIP-seq sample compared to the inputs or controls in the same cell type from within the same study. This data was then ranked from most enriched protein to least enriched protein.

We first looked at the most enriched proteins for each locus and performed a number of quality control checks based on technical and biological expectations. We looked at all of the samples within a study containing an enriched protein, as we observed that problematic samples were often contained within problematic studies. In most cases there was no reason for concern, but examples where enriched-proteins would be excluded include studies with unusually low input/control samples which gave spurious enrichment to all samples in the study, including for proteins which are consistently not enriched in other studies. We also looked at instances of a ‘top hit protein’ in other studies, and noted whether the protein had previously been observed at centromeres in published manuscripts or whether the protein was a new candidate locus-specific protein. We identified the most robust top-hit proteins by focussing on examples that were strongly supported within single studies (i.e. enriched compared to other proteins assayed) and/or across multiple studies.

### REDPOINT validation using published datasets

The BioProject accession number (PRJNA######) for each relevant study is included in the figure legend or supplementary tables associated with each observation.

#### Confirming the centromere localisation of top-hit proteins identified by REDPOINT

To confirm the localisation of the top-hit centromere proteins we repeated the analysis using the complete sequencing run. Each ChIP-seq sample was downloaded using ‘fastq-dump’ from NCBI SRA Tools or directly from the SRA Explorer website (https://sra-explorer.info/). The technical quality of each sample was assessed using FastQC. Each sample was then aligned to the hg38 version of the human genome using Bowtie2 (ver.2.3.0), converted to .bam file using samtools (ver.1.3.1) and to .bed file using BEDTools (ver.2.26). Aligned files were intersected with a table of centromeric regions extracted from the hg38 genome build using the ‘intersect’ function in BEDTools. The total number of aligned reads and the proportion aligned to different repetitive regions, including centromeres, was calculated using custom bash scripts and processed for visualisation in R using ggplot2.

#### Visualising BRD4 as a centromere protein on a T2T genome

To plot BRD4 and other centromere proteins across a complete chromosome we utilised the recently published T2T genome build of human chromosome 8^4^. We followed the same mapping strategy as outlined in the T2T publication^4^, which utilises samtools (ver.1.3.1) and the alignment package bwa (ver.0.7.17)^8^ with the following parameters (bwa mem -k 50 -c 100000 PATH/CHM13.chr8.fasta FILE.fastq > FILE.sam). To convert .sam files to .bam we used samtools sort with the following parameters (samtools sort FILE.sam -o FILE.bam) and samtools index (samtools index FILE.bam). To plot the genomic distribution of BRD4 or other centromeric proteins compared to corresponding input we utilised the bamCompare function in BEDTools (ver.2.30.0) with the following parameters (bamCompare -b1 SAMPLE.bam -b2 INPUT.bam --scaleFactorsMethod SES --operation ratio --binSize 1000 --samFlagExclude 2308 --outFileFormat bedgraph -o NAME.bedGraph). The bedGraph files were uploaded to the UCSC genome browser (https://genome.ucsc.edu/) and the ‘mean’ signal visualised with smoothing set to 10.

#### Identification and validation of JQ1-dependent increase in centromeric BRD4

To validate BRD4-JQ1 as the most prominent hit we included additional ChIP-seq datasets investigating BRD4 following JQ1 treatment which were missed in our annotated REDPOINT data, giving a total of 15 analysed datasets. We followed the same approach outlined above to confirm the centromere localisation using the hg38 and T2T genome. To visualise the fold change in BRD4 localisation following JQ1 treatment on the T2T genome we utilised the bamCompare function in BEDTools (ver.2.30.0) with the following parameters (bamCompare -b1 JQ1_sample.bam -b2 DMSO_sample.bam --scaleFactorsMethod SES --operation log2 --binSize 1000 --samFlagExclude 2308 --outFileFormat bedgraph -o NAME.bedGraph). The bedGraph files were uploaded to the UCSC genome browser (https://genome.ucsc.edu/) and the ‘mean’ signal visualised with smoothing set to 5. To further characterise the effect of JQ1 treatment on centromeric H3K27ac, RNA polymerase II and transcription we analysed ChIP-seq and RNA-seq datasets using the same approaches.

#### Chromosome specificity of BRD4 centromere recruitment by JQ1

We analysed the enrichment of BRD4 following JQ1 at different annotated repeats within the centromere to identify evidence for a central role of CENP-B in recruiting BRD4. We first set out to determine if α-satellite DNA, which is the only centromeric satellite that contains the CENP-B box DNA sequence, is the most enriched repeat class. To do this we aligned matched BRD4-JQ1 and BRD4-DMSO samples to the hg38 genome build, intersected with the table of repetitive regions extracted from hg38, and quantified the percentage of reads aligned to each class of centromeric repeat. Next, we used the same data to focus on the chromosome-specific breakdown of centromeric repeats that recruit BRD4 following JQ1 treatment. We calculated the percentage of centromeric reads aligned to the centromere of each chromosome in each sample. We then determined the ratio of JQ1 compared to DMSO for each centromere in matched JQ1/DMSO pairs to focus on the relative change in BRD4-association across centromeres, independent of centromere length or changes in the total level of BRD4 across all centromeres. Finally, we compared the relative abundance of CENP-B on each centromere to the enrichment of BRD4 following JQ1, using published CENP-B CUT&RUN and BRD4 ChIP-seq data. Enriched chromosomes contain more BRD4 and/or CENP-B than would be expected based on the size of the centromere. The CENP-B data was generated in HeLa, a female cell-line, therefore we cannot assay the Y-chromosome. As negative controls, we compared centromeric CENP-B enrichment to RNA polymerase II JQ1/DMSO treatment and BRD4 following degradation of BRD4 with a PROTAC inhibitor (dBET6/DMSO).

#### Immunoprecipitation mass spectrometry analysis

Processed BRD4 co-immunoprecipitation mass spectrometry (coIP-MS) data was obtained from the supplementary information of two articles^9, 10^. For SUM149/R cells the ratio of JQ1 to DMSO was calculated from the peptide level in the ‘RIME filtered list’ in Supplementary Table S3, and our observations are consistent with data presented in Figure 1E of this study^10^. Data from HEK293T and K562 cells were calculated from Supplementary Table S2G and S2H based on the fold-enrichment of each protein in the pull-down vs control^9^. The ratio of the JQ1 enrichment over the DMSO enrichment was used to identify the proteins most enriched upon JQ1 treatment. Co-IP MS data from this study was also analysed for BRD2 and BRD3 to determine the specificity of the identified drug-dependent protein complexes.

#### Analysis of published CRISPR KO screen in SUM149/R cells

Processed CRISPR KO screen data for SUM149 and JQ1-resistant SUM149R cells in the presence DMSO or JQ1 were obtained from the Supplementary Information of a recent resource article^10^. This data contained the p-values for the enrichment of cells containing each protein-targeting sgRNA compared to control sgRNAs. Enriched sgRNAs have a positive value and highlight proteins whose knock-out promotes cell proliferation, depleted sgRNAs have a negative value and highlight proteins whose knock-out decreases cell proliferation. Our analysis is broadly similar to Figure 1a in the source publication^10^, and uses the same cut-off of p>0.001 for candidate hits but focused on pathways which were not considered or highlighted in the original publication, including proteins involved in centromere function, the spindle assembly checkpoint and sister chromatid cohesion

#### Producing Figures

All data processing and presentation in boxplots, bar charts, line graphs and other forms of graphical analysis was performed in R using standard tools and ggplot2. All immunofluorescence, metaphase spread and western blot images were processed in ImageJ by changing the brightness/contrast only. All figures were assembled in Affinity Designer.

#### Statistical analyses

All statistical analysis was performed in R or Excel and each test and p-value is outlined in the associated legend for each figure.

### Experimental procedures used in validation of REDPOINT results

#### Cell Culture

Human cell lines HEK293T originally obtained from the American Type Culture Collection (ATCC) and HeLa SNAP CENPA (referred to as HeLa cells) were obtained from the Fachinetti lab (Institut Curie). Cells were cultured in DMEM (Gibco #61965-026) supplemented with 10% Foetal Bovine Serum (Capricorn scientific FBS-12A), 1% penicillin/streptomycin (Capricorn scientific PS-B) and sodium pyruvate at a final concentration of 1mM.

#### Bromodomain inhibitor treatment

HEK293T and HeLa cells were treated with 0.25% DMSO (Sigma D8418) or bromodomain inhibitors dissolved in DMSO to give a final concentration of 0.25% DMSO plus inhibitor. Cells were treated for 1 hour or 24 hours, stated in the figure legends, with 500nM JQ1 (Sigma SML0974), 500nM JQ1 (+) or (-) enantiomers (Millipore 500586) or 5μM PFI1 (Torris 4445), a bromodomain inhibitor with a distinct structure and mechanism of action to JQ1 that requires 10x higher concentration for the same reduction in cell growth^11^. In addition, BRD4 degradation was achieved by 24 hours incubation with the PROTAC Inhibitor 500nM MZ1 (provided by Boehringer Ingelheim opnMe portal).

#### Immunofluorescence

HEK293T or HeLa cells were seeded on coverslips in 6-well plates and incubated at least overnight to enable cells to settle and grow on the surface. Cover slips were washed gently once in PBS followed by 7 minutes fixation in 4% PFA, 3x 5-minutes wash in PBS, permeabilization in 0.3% Triton X-100 for 7 minutes and 3x 5-minutes PBS wash. Fixed cells were blocked in 1% BSA in PBS for 30 minutes at room temperature before overnight incubation at 4°C in primary antibodies: α-mouse CENP-A (1:1000 Abcam ab13939 (discontinued) or 1:1500 Enzo ADI-KAM-CC006-E) and α-rabbit BRD4 (1:900 Abcam ab128874). The following day fixed cells were washed 3x 10-minutes in PBS and then incubated 1 hour in secondary antibodies (Alexa Fluor 488, 546 or 647) before 3x 10-minute wash in PBS, 5 minutes incubation in DAPI solution to stain the DNA, 1x PBS wash and mount slide/coverslip with Aqua Poly Mount (Polysciences Inc. 18606-20). Imaging was performed on a Zeiss AiryScan2 LSM900 confocal microscope using the ‘airyscan mode’ and following standard operating procedures^12^.

#### BRD4 co-IP

HEK293T or HeLa cells were split into 1x 10cm dish per IP and allowed to grow overnight before treatment with DMSO, 500nM JQ1 or 5μM PFI1 for 1 hour. We utilised the Active Motif Nuclear Complex Co-IP kit (#54001) following the manufacturer’s instructions with the following variations: we incubated the ‘enzyme shearing cocktail’ for 2 minutes at 37°C, we used the ‘gentle IP buffer’ to wash samples. The immunoprecipitation antibody was α-rabbit BRD4 (Bethyl A301-985-A-M).

#### Western blot analysis

Standard western blot procedures were followed for the quantification of protein abundance in a variety of samples including following BRD4 co-IP. Briefly protein samples were prepared from experimental samples or cell pellets by resuspending in 1x final concentration Laemmli solution, followed by water bath sonication (Diagenode Bioruptor Plus) to reduce sample viscosity and a 5-minute incubation at 95°C to ensure complete protein denaturation. Samples were loaded on a 4-15% gradient gel (Bio-Rad 4561083) and run at 150v for 1 hour (Bio-Rad system). The protein was transferred to PVDF membrane (Amersham 10600022) using semi-dry transfer apparatus using the ‘mixed MW’ setting (Bio-Rad trans blot turbo). Membranes were blocked for 1 hour in 5% powdered milk in TBS-T, followed by overnight primary antibody incubation at 4°C. Primary antibodies used: α-mouse CENP-A (1:1000 Abcam ab13939 (discontinued) or 1:1500 Enzo ADI-KAM-CC006-E), α-rabbit CENP-B (1:500 Abcam ab25734), α-rabbit CENP-C (1:50 Abcam ab193661), β-actin (Sigma A3854) and α-rabbit BRD4 (1:500 Bethyl A301-985-A-M). Following overnight incubation membranes were washed 3x 10-minutes in TBS-T, then incubated in α-mouse-HRP (Sigma A9044) or α-rabbit-HRP (Sigma A0545) in 5% milk for 1 hour at room temperature. Membranes were then washed 3x 10-minutes in TBS-T followed by a brief incubation in ECL reagent. Chemiluminescence was detected using chemiluminescent substrate (Thermo 34095 or 34575).

#### Generating CENP-B CRISPR KO cell lines

CRISPR KO cell lines and matched ‘safe guide’ controls were generated in HEK293T and HeLa cells using lentivirus-mediated gene KO. HEK293T cells were also used to produce the lentivirus. The first step towards generating CENP-B KO cell lines was to establish CAS9 stable cell lines for HEK293T and HeLa cells. Cell lines were transduced with lentiCas9-Blast^13^ (Addgene, #52962) and selected with 20 µg/ml blasticidin to generate stable CAS9-expressing cell lines. sgRNA for CRISPR KO were designed using the Sanjana lab tool (http://guides.sanjanalab.org/) and cloned as previously described^5, 16^. Briefly, sgRNAs were cloned by annealing two DNA oligos and ligating into a BsmB1-digested pLKO1-puro-U6-sgRNA-eGFP. This construct was transformed into Stbl3 bacteria. For virus production 1 million HEK293T cells were transfected with 3 µg of the pLKO1-puro-U6-sgRNA-eGFP plasmid and two helper vectors (2.5 µg psPAX2 and 0.9 µg VSV-G). Packaging cells were transfected using polyethyleneimine (Polysciences, 23966-2) by mixing with DNA in a 3:1 ratio. 48 hours post-transfection viral supernatants were collected, filtered through a 0.45 µm filter and supplemented with 4 µg /ml of polybrene (Sigma) before adding to target cells. Downstream experiments using sgRNA KO were performed on the pool after at least 10 days, or on single-cell clones that were isolated from the pool.

#### Quantifying centromere cohesion defects on metaphase spreads

Metaphase spreads were prepared using a standard method for karyotype analysis using Carnoy’s fixative and dropping the sample onto a microscope slide^15^. This process produces high resolution metaphase chromosome features, but under these fixation conditions centromeric antibodies do not detect their epitope^16^, which prevents fluorescence microscopy. Briefly, 1 well of a 6-well plate per sample was seeded two days prior to metaphase chromosome preparation. 24 hours prior to metaphase chromosome preparation cells were treated with DMSO or BET bromodomain inhibitors (JQ1, MZ1). 2 hours prior to metaphase chromosome preparation cells were treated with 1μg/ml colcemid. Cells were then harvested, combining the media, wash and cells into a single 15ml Falcon tube. To swell cells were resuspended in 5ml 75mM KCl for 10-minutes at room temperature. Cells were pelleted by centrifugation and resuspended in 10ml Carnoy’s fixative (3-parts methanol to 1-part acetic acid) and incubated for 10 minutes at room temperature. This pelleting, resuspending and incubation was repeated twice more and samples were then placed at −20°C at least over-night, and maintained at −20°C for long term storage. To prepare slides of metaphase chromosomes, samples were recovered from −20°C storage, cells pelleted by centrifugation, resuspended in 10ml room temperature Carnoy’s fixative, incubated for 10 minutes at room temperature, pelleted and resuspended in a small amount of Carnoy’s fixative dependent on the pellet size. To drop the slides we took 20 μL of the sample, hydrated the surface of a positively charged microscope slide (Hecht 42409110) with a gentle open mouthed blow, and quickly dropped the sample from a height of approximately 10 cm. Slides were allowed to dry overnight in a dark drawer. The following day slides were rehydrated for 5 minutes in PBS, incubated in DAPI and mounted using Aqua Poly Mount. Imaging was performed on an Zeiss AiryScan2 LSM900 confocal microscope using the ‘airyscan mode’ and following standard operating procedures. Metaphase imaging and analysis was performed blind and the categorisation of centromere defects was based on published observations^17^

#### In vitro reconstitution of the BRD4-JQ1-CENP-B-DNA interaction

To identify whether the BRD4-JQ1-CENP-B-DNA interaction can be reconstituted *in vitro* we purchased or purified each component of the complex. We purchased custom pairs of oligos (Sigma) that can be annealed to form double stranded DNA, and with a 5’ biotin on the forward oligo, for a sequence that has been previously characterised to bind CENP-B *in vitro*^18^ (F primer – GGCCTTCGTTGGAAACGGGATTT) and a shuffled version of the same sequence as a negative control for CENP-B binding (F primer – GTTAGCGATTTGAATTCCGGGGC). Full-length purified CENP-B was purchased from Active Motif (No. 81201) and JQ1 (+) or (-) enantiomers were purchased from Sigma (SML1524 & SML1525).

Sumo tagged BRD4^bromo^ (44-460 aa) was expressed in BL21 *E. coli* after induction with IPTG overnight at 15°C and purified using an ÄKTA pure first with a nickel column (Cytivia HisTrap HP 17524701), followed by a buffer exchange (Cytivia HiTrap Desalting 17140801), Ulp1 cleavage ^19^ and a second round of nickel chromatography to remove the SUMO tag. Protein purity was determined by Coomassie stained SDS PAGE and aliquots flash frozen in liquid nitrogen.

To prepare the magnetic beads, 15μl Dynabeads MyOne Streptavidin T1 beads (Invitrogen 65601) were resuspended in 100μl Binding and Wash (BW) Buffer (5mM Tris-HCl pH7.5, 0.5mM EDTA, 1M NaCl). A magnetic rack was used to separate the beads and the supernatant was discarded. Beads were washed 3x in BW buffer, using the rack to separate the beads and discarding the supernatant each time.

Double stranded DNA was prepared by resuspending complementary oligonucleotides in annealing buffer (5mM Tris-HCl pH6.8, 1mM EDTA, 50mM KCl, 0.01% triton X-100), heating to 95°C for 5 minutes followed by cooling the samples at 1°C per minute until 4°C. 30pmol of dsDNA was resuspended in Binding and Wash (BW) Buffer (5mM Tris-HCl pH7.5, 0.5mM EDTA, 1M NaCl) to a total volume of 100μl. This DNA was added to the pre-washed streptavidin beads and incubated for 30 minutes at 24°C with gentle shaking to immobilise the biotinylated dsDNA on the beads. The beads were then washed one time in BW Buffer and twice in Binding Buffer (20mM HEPES pH7.6, 10% glycerol, 1mM EDTA, 3mM DTT, 0.05% NP40, 150mM NaCl). Following the final wash beads were resuspended in 30μl Binding Buffer containing 2.5 pmol CENP-B, 25 pmol BRD4^bromo^ and 50 pmol (+)-JQ1 or (-)-JQ1. An Input sample 3μl (10%) aliquot was taken and frozen, the remainder of the sample was incubated overnight at 4°C rotating at 20rpm. The following day Binding Buffer was added to a final volume of 300μl, the tube incubated on a magnetic rack, as a Flow Through sample a 30μl (10%) aliquot was taken and frozen, and the remaining supernatant was discarded. The beads were then washed three more times in 300μl Binding Buffer, with a 30μl (10%) aliquot of the Final Wash taken and frozen. After the final wash 30μl 1x Laemmli buffer was added to the beads to give the Complex sample. Each of the Input samples was made up to 30μl in Binding Buffer and 10μl of 4x Laemmli buffer was added to the Input, Flow Through and Final Wash samples. All of the samples were boiled at 95°C for 5 minutes, centrifuged and loaded on 10% SDS PAGE gels for Western Blot. Western blot was performed as outlined previously with primary antibodies to BRD4 (Abcam ab128874) and CENP-B (Abcam ab25734).

## Data and code availability

All data are available in the main text or the supplementary materials. REDPOINT approach utilizes standard analysis tools as described in methods, code available on request.

## Acknowledgements

We would like to acknowledge the bwForCluster Helix and MLS&WISO for computational infrastructure and support. We would like to thank Iain Cheeseman, Andrea Mussachio and Christopher Playfoot for critical feedback on the manuscript and Daniele Fachinetti, Aubry Miller and the Erhardt lab for valuable comments and discussion.

## Funding

Deutsche Forschungsgemeinschaft (DFG) grant EXC81 (CellNetworks) (SE)

Deutsche Forschungsgemeinschaft (DFG) grant GRK2039 (SE)

European Research Council grant ERC-CoG-682496 (cenRNA) (SE)

European Research Council grant grant n°805338 (AB, NSB)

DKFZ Postdoctoral Fellowship (NSB)

Deutsche Forschungsgemeinschaft (DFG) grant INST 35/1597-1 FUGG

Deutsche Forschungsgemeinschaft (DFG) grant INST 35/1134-1FUGG

## Author contribution

Conceptualization: SC, SE

Data curation: SC, NPS

Methodology: SC, NPS, NSB, JB, AMS, VD, SA

Investigation: SC, NSB, JB, AMS, VD

Visualization: SC, NSB, SE

Funding acquisition: SE, AB, NSB

Project administration: SC, SE

Supervision: SC, SE, SA

Writing – original draft: SC, SE

Writing – review & editing: SC, SE, NSB, AB, SA, JB, AMS, VD

## Competing Interests

Authors declare that they have no competing interests.

---

**Extended Data Fig. 1.**
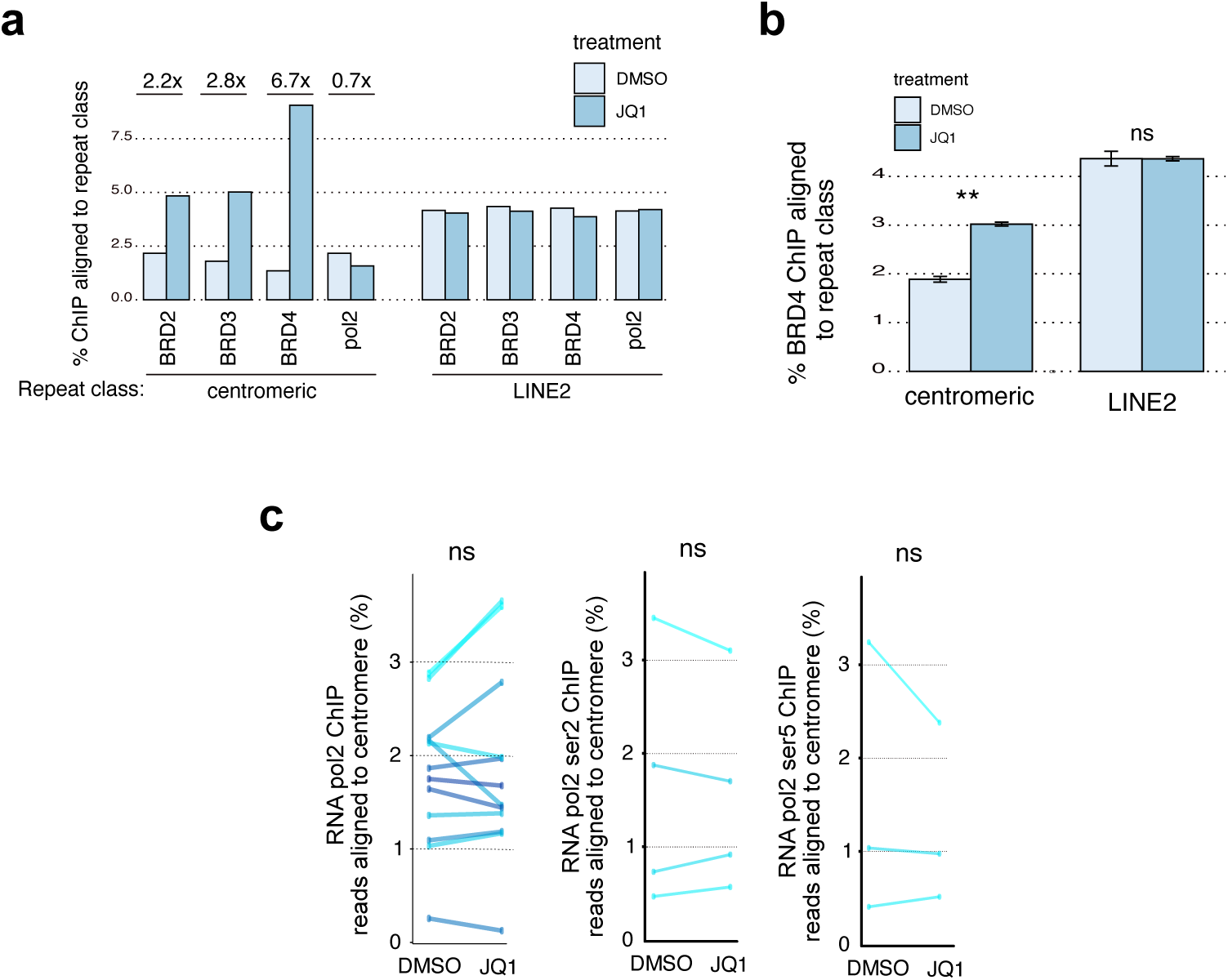
BRD4 is specifically enriched at centromeres following JQ1 treatment, but does not recruit RNA polymerase II. **a**, Centromeric alignment level of BRD2, BRD3, BRD4 and POL2 at centromeres and LINE2 elements following DMSO or JQ1 treatment. Data from: PRJNA253142. **b**, Centromeric alignment level of BRD4 at centromeres and LINE2 elements in a dataset with quadruplicate samples following DMSO or JQ1 treatment. Error bars = standard error of the mean. Statistical analysis t-test (ns ‘not significant’, ** p<0.01). Data from PRJNA482365. **c**, Centromeric alignment of RNA polymerase II following DMSO or JQ1 treatment from multiple studies. Total pol2 – 12 studies, pol2 ser2 – 4 studies, pol2 ser5 – 3 studies. For full data see Supplementary Table 13.

**Extended Data Fig. 2.**
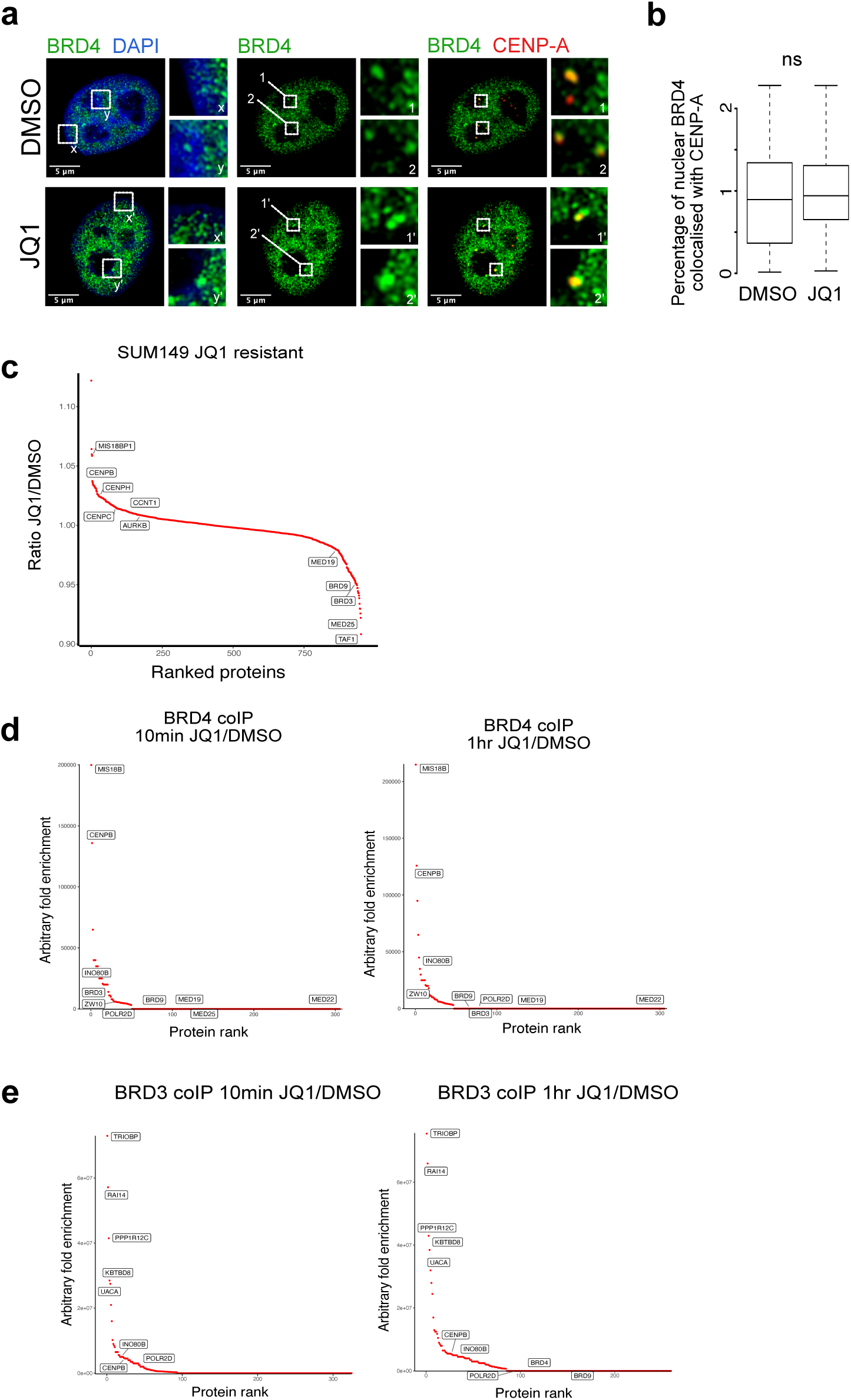
JQ1 stabilises a complex containing BRD4 and centromeric proteins. **a**, Immunofluorescence imaging of BRD4 (green) and CENP-A (red) in HeLa cells following 24 hours DMSO or (±)-JQ1 treatment. BRD4/DAPI inset images highlight the nuclear periphery (x and x’) and the nucleolar periphery (y and y’). BRD4 and BRD4/CENP-A inset images (1, 1’, 2 and 2’) highlight core-centromere locations (CENP-A enriched). **b**, Quantification of the percentage of total nuclear BRD4 signal which co-localises with CENP-A in DMSO or (±)-JQ1 treated HeLa cells. Statistical test is a t-test (n=90 cells for DMSO and 90 cells for JQ1). **c**, BRD4 co-IP MS in SUM149R cells comparing proteins enriched in JQ1 compared to DMSO, with key proteins for centromere identity and BRD4-mediated transcription regulation highlighted. Data from Shu et al. (2020)^28^. **d**, BRD4 co-IP MS in HEK293T cells comparing proteins enriched in JQ1 compared to DMSO after 10 minutes or one-hour inhibitor treatment. Data from Lambert *et al*. 2019. **e**, BRD3 co-IP MS in HEK293T cells comparing proteins enriched in JQ1 compared to DMSO after 10 minutes or one-hour treatment. Data from Lambert *et al*. 2019^29^.

**Extended Data Fig. 3.**
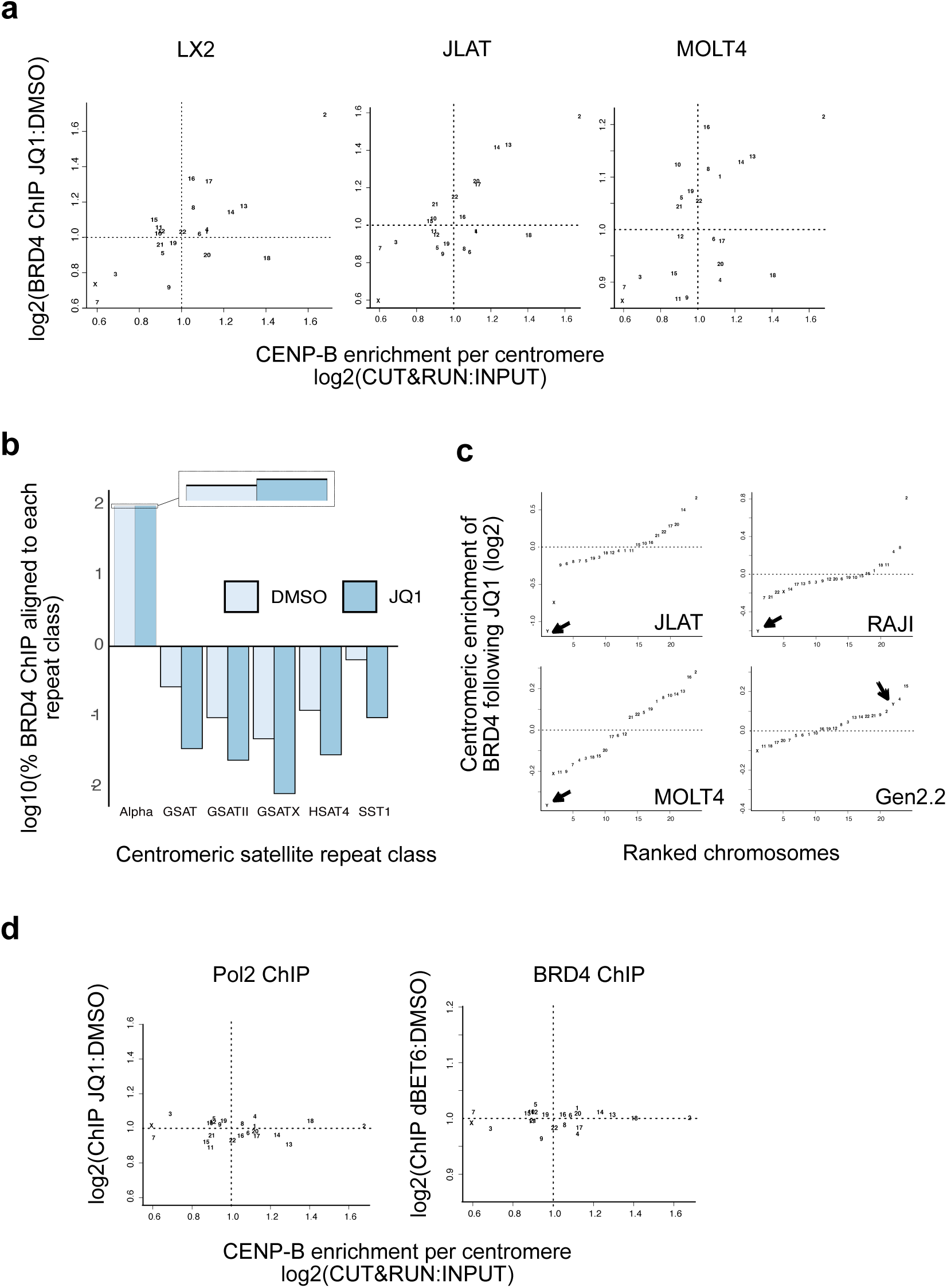
BRD4 is specifically enriched at CENP-B binding sites following JQ1 treatment. **a**, Relationship between CENP-B enrichment at each centromere and BRD4 recruitment following JQ1. CENP-B CUT&RUN data from PRJNA546288. BRD4 data: LX2 PRJNA253142, JLAT PRJNA391180, MOLT4 PRJNA315416. **b**, Quantifying the percentage of reads aligning to each class of centromeric DNA repeat. Whiskers = boxplot standard in R (1.5x IQR). Data in LX2 cells from data from PRJNA253142. **c**, ChIP-seq analysis of centromeres ranked by BRD4 enrichment following JQ1 treatment in male cell lines. Enrichment was calculated by identifying the percentage of centromeric BRD4 at each chromosome in a sample, and then calculating the ratio of JQ1 treated sample compared to DMSO treated sample. The arrow indicates the Y chromosome. Data: JLAT PRJNA391180, RAJI PRJNA306758, MOLT4 PRJNA315416, Gen2.2 PRJNA306431. **d**, Correlation between CENP-B enrichment at each centromere and RNA polymerase II following JQ1 or BRD4 following BRD4 degradation with dBET6. Data: pol2 PRJNA253142, dBET6 PRJNA315416.

**Extended Data Fig. 4.**
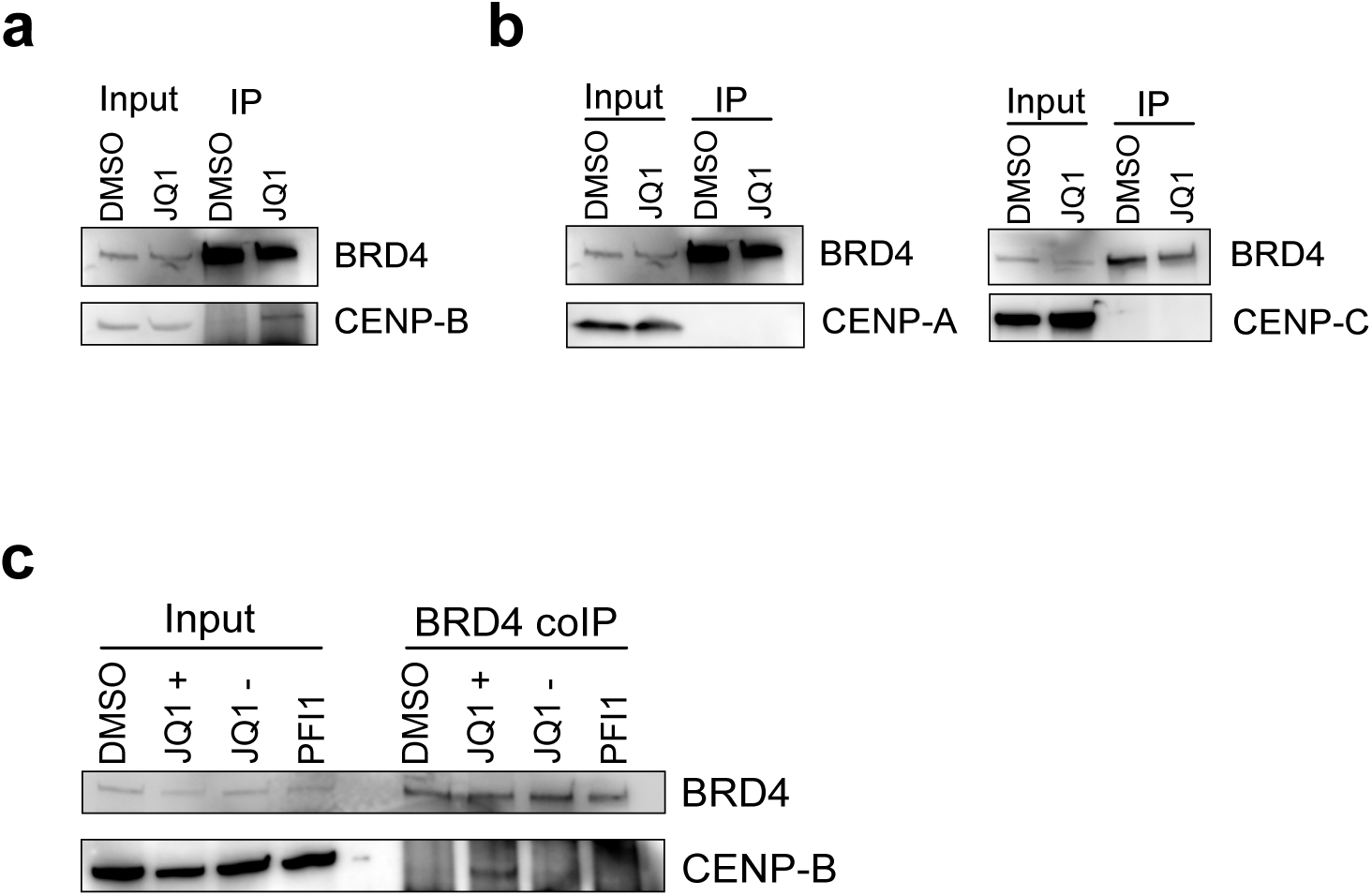
A specific interaction occurs between BRD4 and CENP-B. **a**, Co-IP for BRD4 following DMSO or (±)-JQ1 treatment for 1 hour in HEK293T cells, followed by western blot for BRD4 and CENP-B. **b**, Co-IP for BRD4 following DMSO or (±)-JQ1 treatment for 1 hour in HeLa cells, followed by western blot for BRD4 and CENP-A or CENP-C. **c**, Co-IP for BRD4 following DMSO, (+)-JQ1, (–)-JQ1 or PFI1 treatment for 1 hour in HeLa cells, followed by western blot for BRD4 and CENP-B.

**Extended Data Fig. 5.**
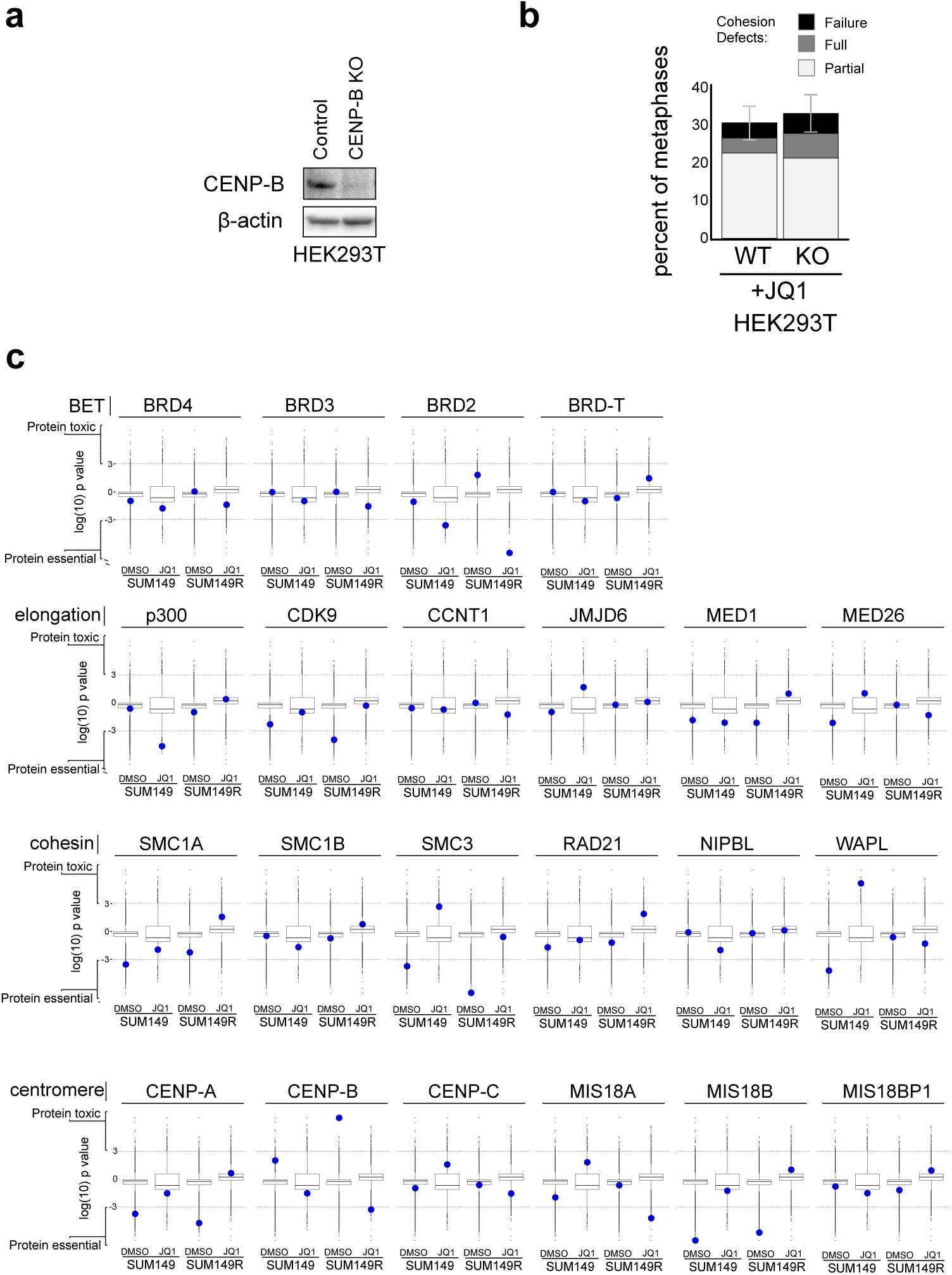
CENP-B has a key role in JQ1-resistance. **a**, Western blot confirming CENP-B depletion in CENP-B KO in HEK293T cells. **b**, Quantification of cohesion defects following 24 hours (±)-JQ1 treatment in HEK293T control (WT) and CENP-B KO cells. Error bars = standard error of the mean. Statistical analysis based on triplicate experiments t-test (ns ‘not significant’). **c**, CRISPR KO screen comparing the effect of different BET bromodomain proteins, transcription elongation proteins, cohesin proteins and centromeric proteins relative to 18,460 other proteins on cellular proliferation in JQ1-sensitive and -resistance SUM149 breast cancer cells. Data from Shu et al. (2020)^28^. Toxic proteins are those in which KO promotes growth, essential proteins are those where KO reduces growth.

## Notes

### Competing Interest Statement

The authors have declared no competing interest.

### Summary of Updates

the revised paper contains improved figures and we removed minor errors from the text.

